# Determining cell fate specification and genetic contribution to cardiac disease risk in hiPSC-derived cardiomyocytes at single cell resolution

**DOI:** 10.1101/229336

**Authors:** Quan H. Nguyen, Samuel W. Lukowski, Han Sheng Chiu, Clayton E. Friedman, Anne Senabouth, Liam Crowhurst, Timothy J.C Bruxmer, Angelika N. Christ, Nathan J. Palpant, Joseph E. Powell

## Abstract

The majority of genetic loci underlying common disease risk act through changing genome regulation, and are routinely linked to expression quantitative trait loci, where gene expression is measured using bulk populations of mature cells. A crucial step that is missing is evidence of variation in the expression of these genes as cells progress from a pluripotent to mature state. This is especially important for cardiovascular disease, as the majority of cardiac cells have limited properties for renewal postneonatal. To investigate the dynamic changes in gene expression across the cardiac lineage, we generated RNA-sequencing data captured from 43,168 single cells progressing through *in vitro* cardiac-directed differentiation from pluripotency. We developed a novel and generalized unsupervised cell clustering approach and a machine learning method for prediction of cell transition. Using these methods, we were able to reconstruct the cell fate choices as cells transition from a pluripotent state to mature cardiomyocytes, uncovering intermediate cell populations that do not progress to maturity, and distinct cell trajectories that terminate in cardiomyocytes that differ in their contractile forces. Second, we identify new gene markers that denote lineage specification and demonstrate a substantial increase in their utility for cell identification over current pluripotent and cardiogenic markers. By integrating results from analysis of the single cell lineage RNA-sequence data with population-based GWAS of cardiovascular disease and cardiac tissue eQTLs, we show that the pathogenicity of disease-associated genes is highly dynamic as cells transition across their developmental lineage, and exhibit variation between cell fate trajectories. Through the integration of single cell RNA-sequence data with population-scale genetic data we have identified genes significantly altered at cell specification events providing insights into a context-dependent role in cardiovascular disease risk. This study provides a valuable data resource focused on *in vitro* cardiomyocyte differentiation to understand cardiac disease coupled with new analytical methods with broad applications to single-cell data.

## Introduction

Genetic effects on early cardiomyocyte development can lead to significant risks of heart defects at birth, and common heart-related health later in life (Bruneau, 2008; Delaughter et al., 2016). While there is some plasticity early in fetal and neonatal development, once cardiomyocytes have reached maturity, the proportion and composition of cellular phenotypes in the adult heart becomes fixed (Porrello, et al., 2011, van Berlo and Molkentin, 2017). As such, many pathological changes in heart development and cellular function are determined during embryogenesis. Population-based studies focusing on cardiac disease, such as coronary artery disease and myocardial infarction, and congenital heart defects have identified 4,517 single nucleotide polymorphisms (*p* < 5x10^-8^) that are significantly associated with disease risk, identifying hundreds of independent genomic loci associated with cardiac diseases (Hoffman and Kaplan, 2002; Bruneau, 2008; Nikpay et al., 2015; Nelson et al., 2017; Howson et al., 2017). The majority of these loci act though changing genome regulation, with their effects observable as expression quantitative trait loci (eQTL), which alter the expression levels disease-associated genes (Koopmann et al., 2014). The discovery of cardiac tissue eQTLs (GTEx Consortium, 2015; Koopmann et al., 2014) is an important step linking cardiovascular disease risk with variation in gene expression. A crucial step that is missing is evidence of variation in the effect of the expression of these genes across the specification of the cardiomyocyte lineage. This problem is addressed at least in part through advances in derivation of human induced pluripotent stem cells (hiPSC) coupled with protocols for cardiac directed differentiation from pluripotency (Eschenhagen et al., 2015, Palpant et al., 2017).

Cardiac tissue engineering by *in vitro* differentiation of hiPSCs is a key approach to developing cell-replacement therapies for heart repair and cardiovascular disease modelling (Garbern and Lee, 2013; van Berlo and Molkentin, 2014; Eschenhagen et al., 2015; Sahara et al., 2015). Current differentiation protocols result in heterogeneous cardiac subtypes such as atrial-, ventricular-, and nodal-like contractile cells as well as non-contractile cell types (He et al., 2003; Ichimura and Shiba, 2017). Transcriptomic studies of cardiomyocyte differentiation based on bulk-sample RNA-sequencing lack the resolution required to determine cell-cell differences, and their contributions to overall phenotype. Critically, the purity of differentiated cardiomyocytes is often assessed by a single marker, for example Troponin T–positive cells (TNNT2+), detected by qPCR or flow cytometry, which mask any underlying heterogeneity (Burridge et al, 2014; Chen et al., 2017). Single cell RNA-sequencing (scRNA-seq) provides an approach to identify discrete cell subpopulations to understand the heterogeneous nature of cellular states defined by differences in transcription. When applied over a time course of differentiation, the scRNA-seq data can be used to map cell subpopulations and identify differentiation trajectories between subpopulations. Importantly, it is also possible to quantitatively estimate the differentiation efficiency (potential for a cell to transition from early to late stages) and heterogeneity (different subpopulations at a single stage) of cells that underlie the differentiation process.

Here, we present results showing the cellular heterogeneity and cell fate choices across *in vitro* cardiomyocyte differentiation, and demonstrate the transcriptional networks governing efficient cardiomyocyte differentiation. We analysed 43,168 single-cell transcriptomes across five differentiation stages during cardiac directed differentiation of human iPSCs using small molecule Wnt modulation. We developed an unsupervised clustering approach, and used this method to identify 15 distinct cell subpopulations and expression signatures with more than 300 new gene markers that distinguish these subpopulations, and show that not all subpopulations progress to the next stages of cell development. Using the transcriptional signature of each subpopulation, we devised a machine learning prediction model, which quantitatively estimates the direction and transition potential for subpopulations within and between different time-points. To determine which genes act as drivers of differentiation, we applied an unsupervised modelling approach, followed by functional and transcriptional network analysis. Finally, we hypothesised that the expression levels of genes associated genetic risk to cardiovascular disease can vary throughout development and, in particular, across lineage specification. We integrated gene expression and GWAS data to identify genes whose expression levels are associated with cardiovascular disease susceptibility, and we show that the expression levels of these genes vary across the cell developmental lineage in a cell fate specific manner. This provides the first evidence for variation in the pathogenicity of a disease loci across a cell developmental lineage.

## Results

### Single cell RNA sequence of cardiac directed differentiation from pluripotency

We induced cardiac directed differentiation using a widely implemented small molecule Wnt modulation protocol (Friedman et al. under-review) and captured cells at days 0, 2, 5, 15 and 30 for scRNA-seq (**Figure 1a**). These five time-points correspond to established cell differentiation states of pluripotency (day 0), germ layer specification (day 2), progenitor (day 5), committed (day 15), and definitive (day 30) cardiac cell types (Paige and Murry et al., 2012; Palpant, et al., 2017). scRNA-Seq data was generated for an average of 4,400 cells from each of two biological replicates at each time-point, with the total dataset comprising of 44,123 cells and 33,020 genes (**Table S1**). Through quality control, we removed 955 cells and 15,302 genes that were expressed in <0.1% of cells (**Table S2**, **Methods**). After filtering, data remained for a total of 43,168 cells and 17,718 genes, with cells sequenced to an average depth of 48,067 mapped reads (**Table S2**, **Figures S1-2**). Filtered expression data were stringently normalised at three levels, per gene, per cell and per sample (**Methods**, **Figure S1**). Following normalisation, we observed no evidence for batch effects based on library preparation, or sequencing run (**Figure S3**). Cells from two biological replicates for each time-point were distributed in close proximity, with no observed differences across days as shown by *t*-distributed Stochastic Neighbour Embedding (*t*-SNE) plots (**Figure S3c**). Similarly, initial hierarchical clustering of the merged data did not show signs of bias across batch labels (**Figure S3b-d**)

**Figure 1.**
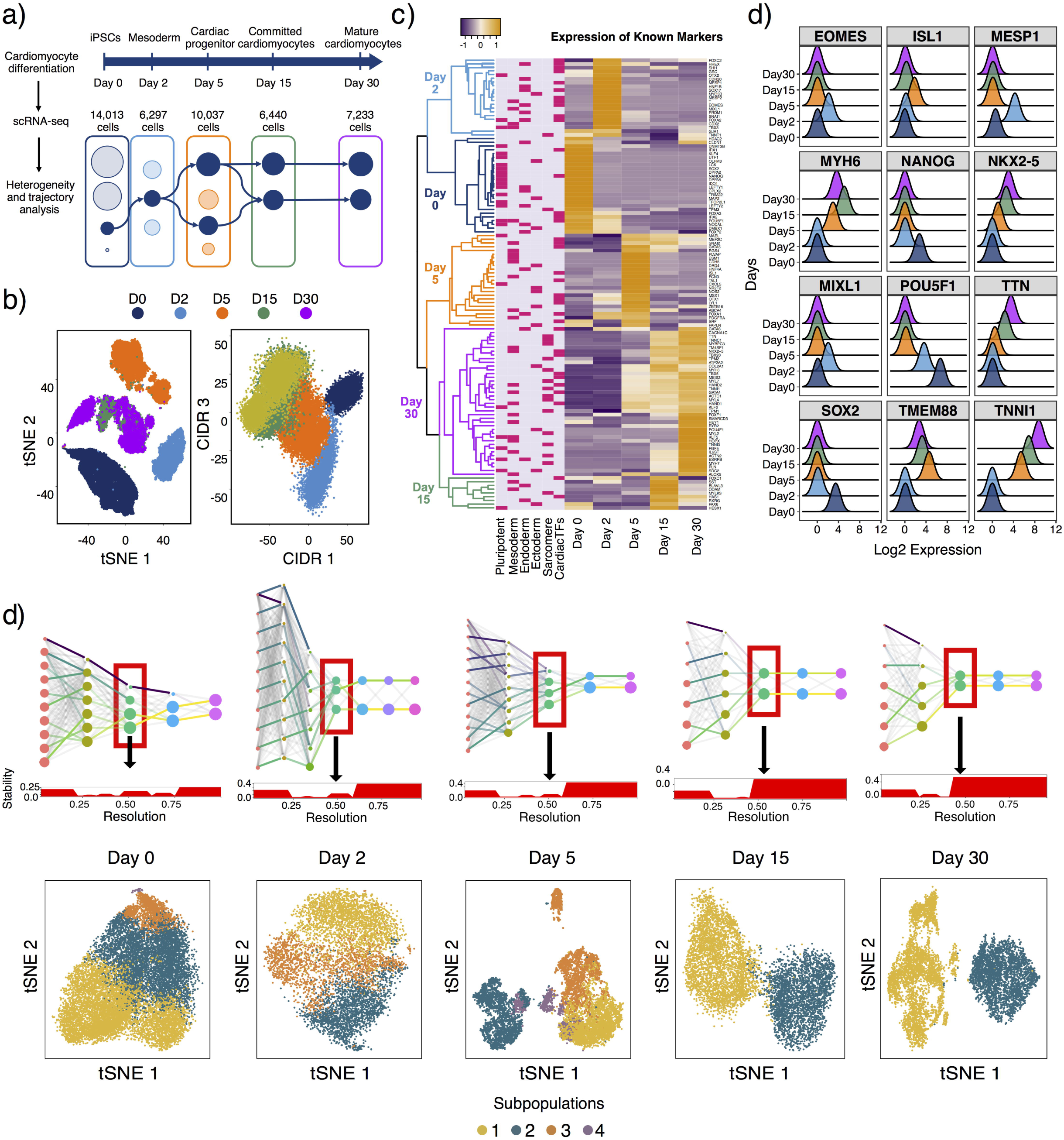
The single cell transcriptomes recapitulate transcriptional heterogeneity at the five cardiac differentiation stages. (a) The study design consisting of differentiation procedure, to single-cell sequencing, and analysis. During the 30-day differentiation process, 44,020 cells were sampled at five differentiation stages: day 0 (pluripotent cells), day 2 (mesoderm), day 5 (cardiac progenitor), day 15 (committed cardiomyocytes) and day 30 (definitive cardiomyocytes). (b) Single-cell data displayed in two-dimensional space by different dimensionality reduction methods, including non-linear embedding with *t*-SNE, and imputed principal components with CIDR. (c) A novel clustering analysis pipeline (Clustering at Optimal REsolution - CORE) to find stable subpopulations. Results are shown for each of the five time-points separately from day 0 to day 30 (right to left). The scRNA data for each day was filtered, normalised, and clustered independently by CORE. The clustering transition can be observed from high resolutions (right) into larger clusters at lower resolutions (left). The horizontal plots show the stability of clustering results. The *t*-SNE plots at the bottom panel display the distribution of cells with cluster colors labelled by the results from the CORE method. Figure S4 contains detailed clustering results for each time point. (d) The expression of 139 core pluripotency and cardiac differentiation markers. The colors represent scaled mean expression of the genes (shown by rows) for each of the five time-points. (e) Expression pattern of known markers for each stage from day 0 to day 30.

Visualizing by *t*-SNE and Clustering from Imputation and Dimensionality Reduction – CIDR (Lin et al., 2017) showed a clear separation of cells transcriptional states corresponding to five discrete time-points (**Figure 1b**). As predicted, cells that were expected to represent pluripotent (day 0), germ layer specification (day 2), progenitor (day5), committed (day 15) and definitive (day 30) cardiac cell types consistently clustered together. To further validate that our scRNA-seq data represented the transcriptional signatures of cell development across the cardiomyocyte lineage, we compiled a list of 139 known markers for pluripotency, germ layer specification, endoderm, ectoderm (Tsankov et al., 2015), and cardiac differentiation (Paige et al., 2012) (**Figure 1d-e**, **Table S3**). The expression levels of these markers strongly confirmed the success of the differentiation process throughout the 30-day time-course (**Figure 1d-e**, **Figure S5**).

Differential expression analysis of genes between cells at days 0 vs. 2, 2 vs. 5, 5 vs. 15, and 15 vs. 30, identified consistent directional fold changes of pluripotent and cardiac developmental markers (**Figures S5-7**, **Tables S4, S5**). The transition from day 0 to 2 shows changes in pluripotency markers such as a reduction in the expression of POU5F1, SOX2, NANOG, and a number of signalling pathways such as WNT (WNT9A, WNT4, DKK1, WNT3A), FGF and FGFR. From day 2 to 5, we observed an increase in the expression of cardiac progenitor genes, including those involved in striated muscle contraction (MYL4, MYL3, ACTN2, TNNI1, TNNT2, MYH6), smooth muscle contraction (MYL7, MYL9, ACTG2, ACTA2, TPM1), and the continuing reduction of pluripotency maintenance genes (NANOG, POU5F1, FOXD3, SOX2). From day 5, cells become committed towards the cardiac lineage, and the expression of genes in the cardiac conduction pathway significantly increase between day 5 to 15 (*p* = 3.4x10^-5^) (**Table S4**). From day 15 to 30, we show that genes associated with cardiomyocyte functions, such as cation transport (RUNX1, SLC22A3) and striated muscle contraction (MYL1, TNNC2) are the most significantly up-regulated (**Table S4**), indicating increasing maturity of the cardiac cells. These results collectively demonstrate that transcriptional variation captured at a single cell level can recapitulate the known functional changes across a cardiogenic differential lineage, but also reveal the substantial limitations of understanding cell fate trajectories from bulk time-point RNA-sequence data.

### Reconstructing the trajectories of heterogeneous cells across the cardiomyocyte lineage

We next aimed to understand cell population diversity represented across each time point during differentiation, and to identify different cell fates of the differentiation lineage. To identify distinct subpopulations of cells, and to identify their trajectories across the lineage, we developed two methods to first classify subpopulations of cells, and subsequently predict their transition to subpopulations at later points within the cardiac lineage.

Normalised expression data were analysed using our novel unsupervised clustering method, <u>C</u>lustering at an <u>O</u>ptimal <u>RE</u>solution (CORE) (**Figure 1c and Figure S4**), to identify subpopulations consisting of transcriptionally similar cells. Using CORE, we identified distinct subpopulations of cells in each of the five differentiation stages (**Figure 1c** and **Figure S4),** which were consistent with clusters separated by the first 2-3 principal components. Our results were consistent with alternative dimensionality reduction approaches, including linear (PCA and MDS), non-linear (*t*-SNE), and imputation approaches (CIDR) (**Figure S4**). Importantly, functional enrichment analysis confirms that these subpopulations are biologically distinct in their differentiation and cell-lineage commitment states (Friedman, et al., under-review, **Figure S6, Figure S7, Table S5**). For pluripotent cells (day 0), we identified four distinct subpopulations representing pluripotency states, including: a core pluripotent (D0:S1 - 6701 cells), a proliferative (D0:S2 - 5614 cells), an early-primed (D0:S3 - 1330 cells), and a late-primed (D0:S4 - 34 cells) subpopulation. This result is consistent with our previous work showing distinct subpopulations of hiPSCs across a pluripotency trajectory (Nguyen et al.). At day 2, we identified a MESP1^+^ mesoderm subpopulation (D2:S2 - 1994 cells) as well as SOX17^+^ definitive endoderm (D2:S1 - 2245 cells) and GSC^+^ mesendoderm (D2:S3 - 1666 cells). Similarly, we observed a diverse cell-type composition at day 5 comprising TNNI1^+^ cardiomyocyte precursors (D5:S1 - 3850 cells), cardiovascular progenitors expressing low levels of cardiac contractile genes TNNI1^+^ cells as well as TAL1^+^ endothelial cells (D5:S3 - 2474 cells), a persistent population of SOX17^+^ definitive endoderm (D5:S2 - 2577 cells) and 10% of cells with no significantly differentially expressed genes (D5:S4 - 996 cells).

Differential expression and gene set enrichment revealed ‘cardiac muscle cell morphogenesis’ as the most significant downregulated pathway in the D5:S2 subpopulation (**Figure S6, Table S5**). This pathway includes known cardiac transcriptional regulators such as HAND1, NKX2-5, WNT5A, FOXC2, and ISL1 (**Figure S6c**), and is in contrast to the upregulation of transcription regulators for endodermal commitment such as FOXA2, SOX17 (**Figure S6b**).

From day 15, as cells become committed within the cardiomyocyte lineage, we observed two major subpopulations, that we identified as non-contractile cells (subpopulation 1: D15:S1 – 3520 cells; D30:S1 – 4038 cells) and contractile cardiomyocytes (subpopulation 2: D15:S2 – 2783 cells; D30:S2 – 3001 cells). Cells identified as cardiomyocytes in D15:S2 and D30:S2 expressed more definitive cardiomyocyte markers, specifically transcription regulators (NKX2-5, FOXC2, IRX4) and cardiac structural genes (ACTC1, MYH7, MYH6, MYL4, TTN, TNNT2, TNNI1, TNNC1) (**Figure S7d, e; Table S5**, and **Figures 2 and 4** in Friedman et al., under-review), whereas the non-contractile cells in D15:S1 and D30:S1 expressed markers consistent with an outflow tract-like cell (THY1, KDR PITX2, PDGFRB, BMP4, FGF10). Extensive analysis of cell subpopulations and transcriptional comparison to in vivo cell types is provided in our companion study (Friedman et al, under review).

**Figure 2.**
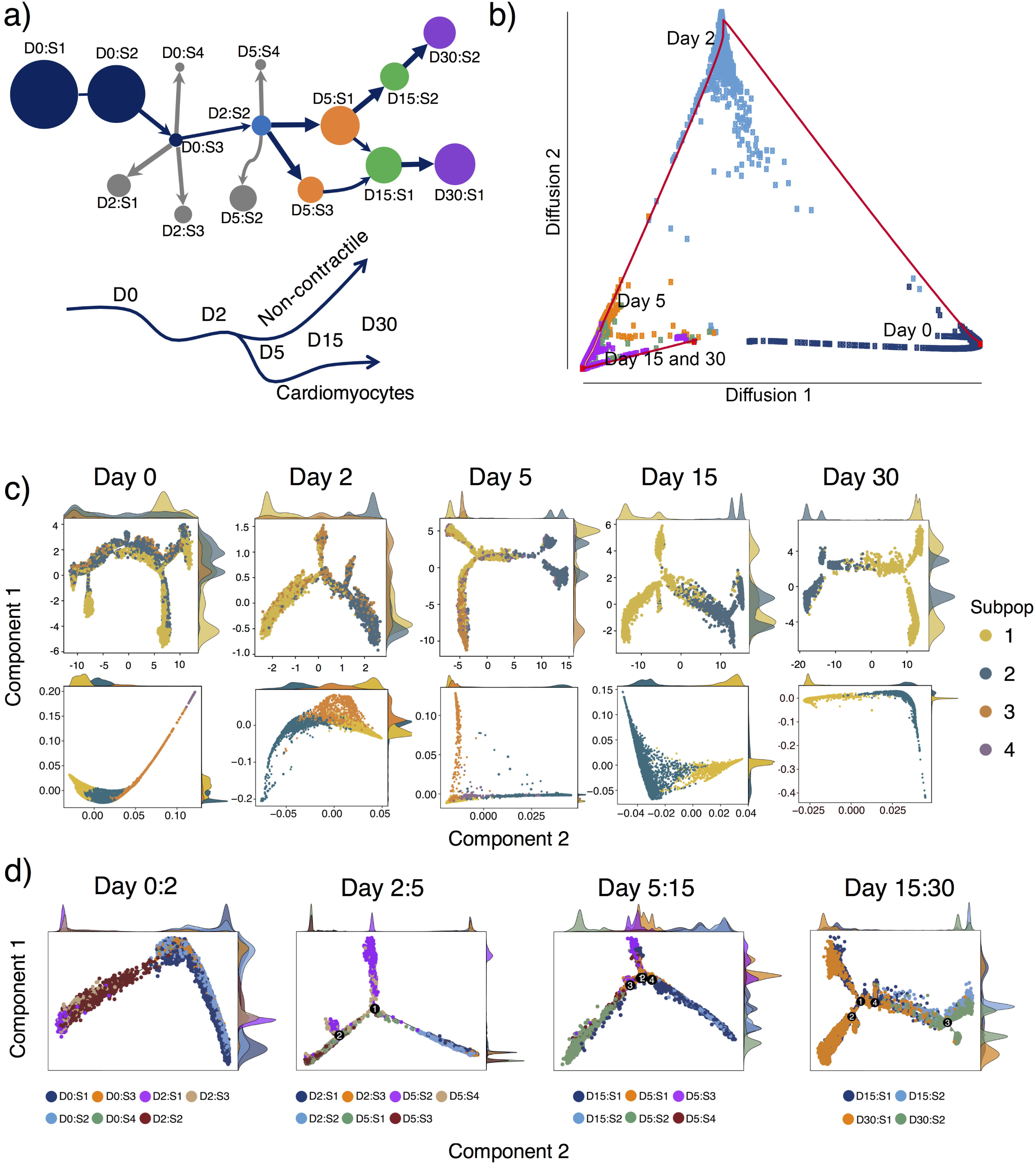
Decomposing the differentiation pathways and subpopulations from hiPSC to mature cardiomyocytes. (a) Differentiation trajectories from day 0 to day 30, predicted by the scGPS. Each of the 15 distinct subpopulations identified by CORE are presented as a circle, with the area proportional to the number of cells, and colors demote sequencing time points. The arrow widths are proportional to the transition scores (0 - 100%, **Table S7**). The bifurcation event from progenitor cells on day 5 to two branches on day 15 and day 30 is identified by the transition scores. (b) Cell trajectory based on diffusion pseudotime. The pseudo trajectory starts from day 0 and follows the temporal order to days 2, 5, 15 and 30. (c) Within time-point transition between subpopulations. Differentiation pseudotime was calculated by Monocle2 (top panel) and by Diffusion (bottom panels) for cells in days 0, 2, 5, 15, and 30 (right to left). Two cells that are at the similar stage/state are at closer proximity in the trajectory. The density on the axes show how subpopulations intersect. The same figures with pseudotime scale are shown in the supplementary **Figure S12**. (d) Monocle2 analysis of the transition between two continuous days, including: day 0 to 2, day 2 to 5, day 5 to 15 and day 15 to 30.

To confirm that clustering was not an artefact of differences in cell cycle, we estimated the probability of a cell being in G1, S, or G2M stages of the cell cycle (**Table S6**). In total, 36,419 cells in G1, 4,528 cells in S phase and 1806 cells in the G2M phase. The majority of the cells (>75%) in each cluster are in the G1 phase **(Figure S9**).

To facilitate public access and interpretation of time-course scRNA-seq data, we have created a publicly accessible database and analysis server (**Figure S8**), which can be accessed here: http://computationalgenomics.com.au/shiny/hipsc2cm/.

### A novel differentiation trajectory analysis framework

To study cell transition and fate choices of the different distinct subpopulations of cells, we developed a method to predict the transition of cells in one subpopulation to cells in a subsequent subpopulation. This method is called <u>s</u>ingle <u>c</u>ell <u>G</u>lobal fate <u>P</u>otential of <u>S</u>ubpopulations (scGPS), which trains a regularized logistic classifier to estimate the probability of cells transitioning to another subpopulation. We applied this between subpopulations of cells within a time point, and between adjacent time points (**Figure 2a, Figure S10 and Table S7**). Cell lineage commitment revealed by trajectory or pseudotime analysis often focuses on continuous transition between neighbouring cells (Haghverdi, 2016; Qiu et al., 2017), but not between discrete subpopulations, making fate choices difficult to identify especially in time course single cell data that lack continuity of transcriptional states between cells through differentiation. The scGPS method addresses this problem by estimating a probability that cells in one subpopulation are able to transition to another subpopulation. We call this probability a transition score. The combination of CORE and scGPS methods form a framework that can be applied for any unknown (sub)population of cells to classify cells into subpopulations, to objectively identify gene markers and to estimate transition scores between the subpopulations (**Figures S10 and S11, Table S7**). The prediction of transitions cell subpopulations by scGPS is consistent with Monocle2 (Qiu et al, 2017) and Diffusion pseudotime (PT) (Haghverdi et al., 2016) (**Figure 2**). However, scGPS has multiple advantages over these methods. In particular, scGPS quantitatively estimates the transitioning potential of each subpopulations within and across a time course. Compared to existing methods, scGPS does not assume a linear continuum trajectory in the total population, and can statistically predict the transition between any pair of subpopulations. Importantly, scGPS can be applied in an unsupervised manner to both discover novel trajectories and new markers. Alternatively, a transition score can be calculated using a gene list of known markers.

To illustrate the value in using scGPS to identify novel genes governing cell fate decisions, we estimate the transition scores for all subpopulations across the time-series using the unsupervised approach, and also the list of 139 cardiac lineage markers (**Table S3**). In all cases, the transition score, and deviance explained were consistently higher (0.2-8.6%, and 6-44% respectively) when scGPS was trained using unsupervised method, compared to known the pluripotency and cardiogenic markers (**Figure S11, Table S8**). These results provide evidence that our analysis of single cell transcription profiles can identify novel drivers of fate choice and cell differentiation consistently across the cardiomyocyte lineage. For example, scGPS identifies PAX2 with the highest coefficient for the D5:S1 subpopulation, which is the most cardiogenic subpopulation giving raise to day 15 cells (see below). PAX2 has not been reported to be involved in cardiomyocyte differentiation, but is highly expressed heart tissue, suggestive of a gene important in early heart morphogenesis. As expected, scGPS also identified genes known to be involved in cardiomyocyte development, with positive coefficients estimated for TNNC1, NKX2-5, MYL7, and MYH6 in the derivation of D30:S2 cardiomyocytes. In contrast, PDGFRB and HOXA3 had a negative coefficient for D30:S2, and are involved in non-cardiomyocyte fates. Our results demonstrate that scGPS is able to identify both known markers of cell identity, and reveal extensive new statistical evidence for novel genes underlying cell subtype specification during cardiac differentiation.

Using scGPS, we were also able to identify transitions between subpopulations within a given time point (**Figure 2c**). For example, for cell subpopulations identified at day 0, we identified a transition from pluripotency to late primed cells consistent to our previous work using hiPSCs (Nguyen et al.). Over 70% of the cells in the D0:S1 were capable of transitioning into the D0:S2, while D0:S2 and D0:S3 had the highest potential transition scores (~80%) to D0:S3 and D0:S4 respectively (**Figure 2a, c**, **Table S7**). The scGPS data for day 0 is in agreement with the observed trajectories identified from the Diffusion prediction method (Haghverdi et al., 2015, 2016) (**Figure 2c**), and Monocle2 (Qiu et al., 2017) (**Figure 2c**, **Figures S10 and Table S7**).

### Cell fate of subpopulations across differentiation stages identified using scGPS

For cells sequenced at day 2, the D2:S2 (mesoderm) subpopulation, characterised by high expression of the cardiac mesoderm gene MESP1, had the highest potential to transition into cells in D5:S1 (cardiomyocyte precursors), D5:S3 (cardiovascular progenitors) and D5:S4 (intermediate) subpopulations in day 5 (**Figure 2a**, **Table S7**). Analysis of the transition between day 5 and 15 demonstrates the progression of the cardiomyocyte precursors (D5:S1) and cardiovascular progenitors (D5:S3) (**Figure 2**) and the termination of the definitive endoderm (D5:S2) in the cardiomyocyte differentiation trajectory. In contrast, subpopulations D5:S1 and D5:S3 both transition into the non-contractile cardiovascular subpopulation (D15:S1). Interestingly, from day 15 to day 30, we identified a consistent overlap between two the subpopulations of non-contractile cardiovascular subpopulations (D15:S1 vs. D30:S1) and contractile committed/definitive cardiomyocytes (D15:S2 vs. D30:S2), providing evidence of bifurcation at day 5 into two distinct lineage branches (**Figure 2d**). One of these branches forms non-contractile, fibroblast-like cardiac derivatives (with high expression of outflow tract cell markers such as THY1, PITX2, KDR, PBX2 and BMP4 and low NKX2-5 expression), and the other branch forms a mature, contractile heart muscle cardiomyocyte subpopulation with strong expression of contractile cardiomyocyte markers such as TNNI1, TTN, MYL2. *In vivo* functional analysis of the two subpopulations showed a reduction in contractile force of the D30:S1 compared to D30:S2 subpopulations (*p* = 0.0009) (Friedman *et al.,* under review). Moreover, the scGPS results for subpopulations in days 15 and 30 predicted that over 90% of cells in D30:S1 and D30:S2 are direct transitions of cells in D15:S1 and D15:S2 respectively (**Figure 2a**, **Figure S10**). Additional validation of the scGPS prediction algorithm demonstrated the reproducibility of known, *in vivo* developmental lineage choices (Friedman et al., under review). Finally, we separately normalised data consisting of pairs of time points, from day 0 to 2, 2 to 5, 5 to 15, and 15 to 30, and implemented Monocle2 and Diffusion pseudotime methods to reconstruct pseudotime differentiation trees between these time-points. The results strongly support our conclusions regarding cell fate trajectories (**Figure 2d, Figure S12**).

### Co-expressed genes are dynamically regulated across the cardiomyocyte lineage

We hypothesised that during differentiation, gene expression was dynamically regulated for co-expressed modules governing gene networks from pluripotency to committed cardiac fates. Gene regulatory circuits, which contain a large number of genes that are mutually inhibited or stimulated, have been shown to underlie cell fate decision in systems, such as the differentiation from bone marrow-derived cell line to erythroid and myelomonocytic lineages (Huang et al., 2007), and from mouse embryonic stem cells to mesendodermal cells (Shu et al., 2013). In line with these studies, cellular reprogramming from a defined intermediate population in mouse, was shown to involve nine gene categories dynamically regulated by two distinct transcriptional waves (Polo et al., 2012). These modules consist of co-ordinately regulated genes, including master regulators (e.g. transcription factors) and effector genes (e.g. structural genes or genes related to physiological activities), enabling subsequent identification of pathways potentially driving differentiation.

We initially identified a total of 483 genes whose expression levels were significantly (*p* < 0.05 / 17,718) associated with polynomial model of the differential time-course (**Figure 3a, Tables S9, S10, S11**). We stratified these genes into co-expression modules, based on topological analysis of gene expression co-regulation (**Figure 3b**). This identified two distinct modules. The first, which we termed the ‘early’ module, contains 367 genes, whose expression levels decrease as cells differentiate into mature cardiac fates. The second module, termed ‘late’, contains 116 genes that are up-regulated as the cells exit from pluripotency into differentiated states (**Figure 3, Tables S9, S10, S11**). Interestingly, the between-module gene correlations are in opposite sign directions, suggesting independent regulatory networks **(Figure 3c, Table S11)**. Examples of hub genes in these two modules include a pluripotency marker, POU5F1 (early module), and a cardiac conduction marker, TPM1 (late module). From day 0 to day 30, when cells progressed from pluripotency into the cardiac lineage, POU5F1 expression decreased, while TPM1 expression increased (**Figure 3c**).

**Figure 3.**
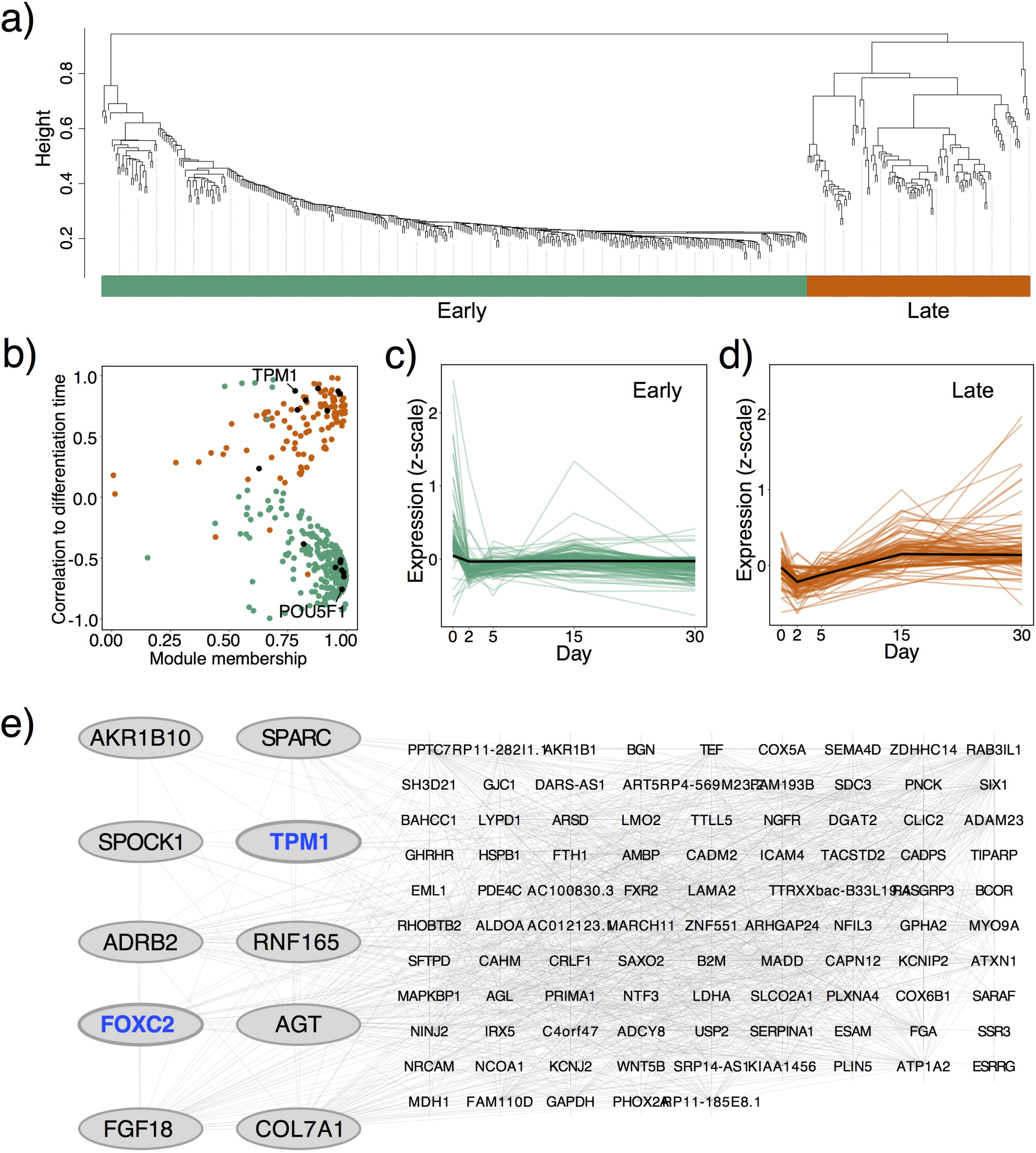
Dynamic changes in gene expression across the differentiation lineage. (a) Gene co-regulation analysis of the 483 genes whose expression levels significantly (*p* < 2.8x10^-6^) change across the lineage. We identified two co-regulated modules, which we have denoted as early (higher expression on days 0, 2 and 5) and late (higher expression on days 15 and 30). (b) The module membership and correlation differentiation time shows little overlap in the modules. Genes with both a high module membership and correlation are suggestive of being key genes within the module networks. Genes with a significant association to cardiovascular diseases from the SMR analysis are shown as black points. (c) and (d) show the standardised expression of each gene in real time-scale (from day 0 to 30). The black lines are the mean trend lines, showing the decreasing (early) or increasing (late) gene expression during the differentiation time-course. (e) Transcription factor network analysis for the co-regulated late module. We identified FOXC2 as the key gene, both in the number of connections and in the correlation to the differentiation time-course. TPM1 has high module membership and correlation to differentiation time, and also has cardiac tissue eQTLs (Koopman et al., 2015).

To identify the mechanisms controlling the co-regulation of expression, we tested for enrichment of transcription factor (TF) binding targets for genes within the early and late modules (**Figure S13**). The early module contains gene targets enriched for 16 transcription factors (Normalised Enrichment Score >= 3) (**Table S11**). In particular, ZFP42 is predicted to target 62 genes in the early module, and the critical role of ZFP42 in governing the exit from pluripotency and germ layer commitment has been confirmed *in vitro* and *in vivo* (Kalkan et al., 2017). Similarly, we identified 14 independent TFs that are significantly enriched for gene targets in the late module (NES >= 3) (**Table S11**), with the most highly enriched being MEF2A (Myocyte Enhancer Factor 2A), a core cardiac transcription regulator (Desjardins and Naya, 2016) that binds to 55 of the 116 genes.

While our initial data provide strong evidence for co-regulated early and late gene networks identified using a polynomial model to analyse the time course differentiation data, we next utilized this approach to further understand cell fate diversification of subpopulations. To gain insights into genetic regulators of subpopulation lineage trajectories, we conducted the gene-model analysis with trajectories identified by scGPS (**Figure 2a)**. The two definitive cardiac fate trajectories identified at day 15 of differentiation differ from the bifurcation in day 5 (**Figure 2a**). Our annotation analysis based on cardiac developmental biology (Friedman et al., under review) suggests that trajectory 2 gives rise to contractile cardiomyocytes, while trajectory 1 generates non-contractile cells with an outflow tract (OFT)-like transcriptional signature.

We found a strong distinction between the two trajectories in the gene-model analysis. Trajectory 2 had more genes whose expression levels significantly change over the time-series (498 genes vs. 136 genes), and more trajectory-specific genes (110 genes vs 31 genes) (**Table S12**). Trajectory 2 contains significant genes enriched for cardiomyocyte differentiation (**Table S12**), and iRegulon analysis of genes in trajectory 2 identified key cardiomyocyte regulators, including MEF2D, MEF2C and MEF2A, as the highest ranked transcription factors (**Table S11**, **Figure S13**). Importantly, we observed two key OFT markers, namely PBX1 (Pre-B cell leukemia transcription homeobox) and SOX17 (SRY-box 17), that were enriched for transcription factors for trajectory 1 genes (Verzi et al., 2005; Buckingham et al., 2005; Chang et al., 2008). These results further support the biological annotation of the pathway to outflow tract (trajectory 1) and to definitive cardiomyocytes (trajectory 2) and provide insights into co-regulated transcriptional networks underlying cell fate diversification during differentiation.

### Comparison of gene expression in single cardiomyocytes with bulk RNA-seq of postnatal and adult heart

*In vitro* differentiated cardiomyocytes are transcriptionally distinct from postnatal and adult heart cells, particularly in terms of regeneration potential (Garbern and Lee, 2013; Sahara et al., 2015). In the context of translational functions of stem cell-derived cardiomyocytes, it is crucial to understand the differences in gene expression programs that are active in differentiating cardiomyocytes and the effect they have on downstream processes. We sought to compare the expression profiles of single cells at five stages of *in vitro* cardiomyocyte differentiation with expression in post-natal to adult heart tissues.

We identified a high correlation between prenatal and adult heart expression levels (Spearman’s *ρ* = 0.8-0.9), and a high correlation between the differentiation days of the individual single cell cardiac directed differentiation samples (*ρ* = 0.7-0.9) (**Figure 4e**). Importantly, we show that the correlations between mature cardiomyocytes (day 30) and post-natal heart (*ρ* = 0.6) and healthy adult heart (*ρ* = 0.5-0.6) is higher than that between the adult heart tissue (age 34) and the day 0 pluripotent iPSCs (*ρ* = 0.4) and the day 2 mesoderm cells (*ρ* = 0.4). The significant correlation between early/late *in vitro* differentiation of iPSCs to cardiomyocytes with post-natal and adult heart indicate that the transcriptomes of differentiated cardiomyocytes reliably reflect human cardiomyocyte gene expression however further studies are required to derive the subpopulation diversity of the adult heart from pluripotent stem cells.

**Figure 4.**
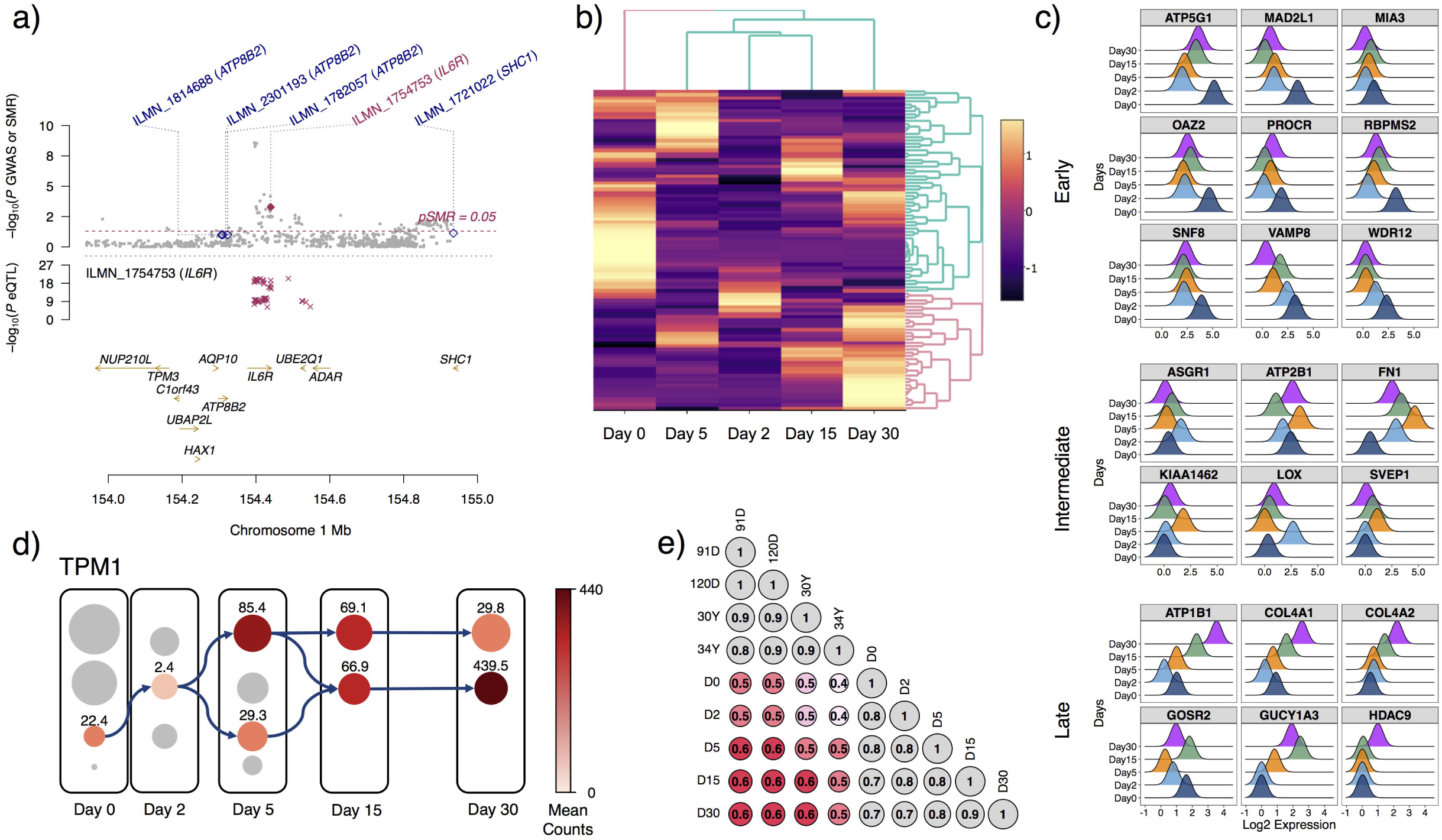
Cardiac disease genes identified from integrating single-cell data with large-scale population genetic data. (a) Combining GWAS and eQTL data revealed IL6R as a potential causal gene of cardiac disease. (b) 95 coronary artery disease (CAD) related genes (CAD genes) identified from GWAS were highly expressed at specific differentiation stages. (c) Expression of 21 CAD genes with significantly altered expression patterns between time-points. Genes were grouped as highly expressed during early, intermediate and late differentiation. (d) Fate specific (shown by two differentiation trajectories) expression levels of TPM1, whose expression levels are significantly associated with an increase in cardiovascular disease risk. (e) Correlation of scRNA-seq expression levels (from pluripotent to differentiated cells) with bulk RNA-seq expression levels in ENCODE heart foetal and adult tissues.

### Integration of population-level disease genomics results with single cell cardiomyocyte lineage data

A recent study of risk loci for coronary artery disease (CAD) (Howson et al 2017) highlighted 95 candidate causal genes with functional data that are associated with arterial wall mechanisms. We hypothesised that the expression of CAD genes could vary across the differentiation time-course, and between the two cell fate trajectories. Using the scRNA-seq expression data for the 95 CAD genes, we selected genes with the highest overall expression levels (**Figure 4b**), and grouped them as pluripotent (Day 0), mesoderm (Days 2 and 5), and cardiomyocyte (Days 15 and 30) stages. We found that all 95 genes have significant changes in expression across days (**Figure 4c, Tables S15-** 16), suggesting that any significant eQTL underlying the expression levels of these genes may have a variable effect across the cardiac developmental lineage.

Genetic variants that contribute to common disease risk are known to predominantly act through changes in gene expression. Importantly, to date, most of this knowledge comes from analysis of gene expression measured from bulk tissue, containing predominately mature cells, meaning gene expression changes that occur across a cell developmental lineage are ignored. Our study presents the first opportunity to investigate how the expression levels of cardiovascular disease genes vary across the developmental lineage, and between different cell fates. We first identified cardiovascular disease genes using Summary Mendelian Randomization (SMR) to test for shared causal loci between 2,962,408 cardiovascular GWAS SNPs and 15,248,720 eQTL SNPs identified from expression levels measured using bulk RNA-seq in cardiac tissue. Our analysis yielded a total of 226 significant genes whose expression levels are associated with cardiovascular disease risk because of a shared causal variant (**Table 1**). For example, in **Figure 4a**, we provide an example of a gene candidate identified from the SMR analysis, with significant cardiac-related eQTLs acting in a causal regulatory role, IL6R, a well-studied cardiac disease-causing locus (Sarwar et al., 2012) is shown in **Figure 4a**.

**Table 1.** Results from the Summary Mendelian Randomisation (SMR) analysis of cardiac disease GWAS and eQTL data.

|  | Genes | eQTL SNPs<br>(MAF >=0.01) | GWAS SNPs<br>(MAF >=0.01) | SNPs retained from<br>ref. genotype | Total SMR<br>Genes | Sig. SMR<br>Genes | Unique SMR<br>Genes | Unique SMR<br>SNPs |
| --- | --- | --- | --- | --- | --- | --- | --- | --- |
| Left Ventricle<br>(GTEx) | 21,709 | 6,992,633 | 2,962,408 | 2,170,851 | 1907 | 53 | 53 | 52 |
| Atrial Appendage<br>(GTEx) | 22,688 | 6,837,657 | 2,962,408 | 2,166,412 | 1607 | 40 | 40 | 39 |
| Left Ventricle<br>(Koopmann) | 429 | 5628 | 2,962,408 | 3333 | 118 | 4 | 2 | 2 |
| Peripheral Blood<br>(CAGE) | 25,422 | 1,412,802 | 2,962,408 | 393,097 | 7212 | 175 | 159 | 161 |

We subsequently investigated the changes in gene expression for the 226 genes across the cardiomyocyte differentiation time course based on: (i) membership to early and late regulatory modules; (ii) differential expression between cells at differentiation time points; (iii) between subpopulations for specific time points. Our results identified 15 genes whose expression levels are associated with cardiovascular disease and that change significantly across the cardiomyocyte developmental lineage (**Tables S13, S14**). Of these, we observed an overlap of 11 genes with the ‘early’ network module (ATP5G1, CCDC15, DNAJB6, FLVCR2, IFI16, NFE2L3, POU5F1, RFC4, RTN4, SMARCB1 and SUMO2), and four with ‘late’ network module (ADRB2, GAPDH, MADD and PPTC7) (**Table S14**). Notably, the ‘early’ module genes included POUF51 (OCT4), FLVCR2, a calcium transporter, and DNAJB6, a known cardiomyopathy susceptibility gene (Ding et al, 2016). To more clearly describe the value of integrating SMR data, an example is provided using DNAJB6, which is associated with cardiovascular disease. An eQTL analysis revealed that the SNP rs3802096[C/T] is significantly associated with changes in the expression level of DNAJB6 (beta = -0.495, *p* = 1.70 x 10^-74^) and SMR provided evidence the expression levels were associated with cardiovascular disease (adjusted *p* = 0.0332). In our scRNA-seq data, we observed a 2.71-fold decrease in expression between day 0 and day 30, suggesting allelic effects on expression will have the greatest effect early in the cardiac cell developmental lineage. Moreover, several genes that we identified as important in cardiac regulatory networks (POU5F1, ATP5G1, ADRB2 and SMARCB1), were recapitulated in the SMR analysis, which provides evidence that the genetic control of cell fate choices can impact on cardiovascular disease risk. However, further studies using animal models coupled with manipulation of gene dosage is required to confirm a developmental role for these cardiovascular disease risk loci.

Finally, we investigated changes in the expression levels across the cardiac developmental lineage for 406 genes with a cardiac tissue eQTL (Koopmann et al., 2014). Of these genes, 13 had significant changes in their expression across the lineage, 10 in the ‘early’ and 3 in the ‘late’ module (**Table S13**). One interesting example is tropomyosin 1 (TPM1), which is regulated by the rs4479177 SNP in Koopmann et al. (left ventricle *p* = 2.8x10^-6^) and GTEx (left ventricle *p* = 1.0x10^-4^; atrial appendage *p* = 1.72 x 10^-3^). In our single cell data, TPM1 is highly up regulated in D15:S2 (2.2-fold increase compared to D15:S1), which represents the most developmentally mature cardiomyocytes, and D30:S2 (9.2-fold increase compared to D30:S1) (**Figure 4d**, **Table S13**). Importantly, TPM1 is the key hub gene of the late module (**Figure 3f**).

Collectively, these results demonstrate that by integrating population-level data on the genetic control for common diseases, with scRNA-seq for cell lineages, we are able to gain valuable knowledge on how the pathogenicity of a loci can vary as cells develop from pluripotent to mature states.

## Discussion

The development of cardiac stem cells for therapeutic or translational applications requires an understanding of mechanisms underlying hiPSC differentiation into committed and mature cell (sub)populations (Garbern and Lee, 2013; Sahara et al., 2015). Here, we decipher important aspects of this process, with particular emphasis on the proportion of transitioning cells, the composition of differentiated populations, and molecular mechanisms with regulatory genes and networks driving heterogeneity during cell fate determination. We identified differentiation trajectories underlying cardiac directed differentiation by Wnt modulation and quantitatively characterized the heterogeneity among subpopulations from pluripotency to lineage commitment and maturation.

Clustering cells into subpopulations is a crucial step that determines downstream analysis in singlecell workflow. Often, clustering algorithms require modelling and parameter optimisation, which are computationally intensive (Lin et al, 2016). Computationally fast scRNA clustering methods, such as K-means clustering and SC3 (spectral clustering) (Kiselev et al., 2017) rely on *a priori* assignment of the number of clusters, and thus are less suitable when novel, rare, or complex biological subpopulations exist. We developed the CORE algorithm with an unsupervised approach that does not require user defined-parameters, and provides statistical confidence for the optimal number of clusters that can be identified from a scRNA-seq dataset. CORE is computationally fast, and designed to identify complex clustering patterns including nested clusters, detect outlier cells, and determine a stable, optimal number of clusters. Here we show that CORE recapitulates the diversity of developmental cell populations involved in governing cardiac development *in vivo* and *in vitro*.

Cell lineage commitment revealed by trajectory analysis is often achieved by reconstructing pseudotime differentiation paths from the undifferentiated (root) to differentiated branches (Haghverdi, 2016; Qiu et al., 2017). These approaches are able to estimate transition probability between cells, but not between distinct subpopulations of cells. Specifically, these methods arrange cells into continuous differentiation paths from a root to terminal branches, but do not estimate transition probability between cells or subpopulations that are not neighbouring in the differentiation trees. Our scGPS method decomposes cells into discrete subpopulations and estimates a transition probability between subpopulations. Thus, scGPS addresses the question of how to predict subpopulation transitions. Importantly, the lineage trajectories identified by scGPS fit with known developmental trajectories of cardiac development from pluripotency, but in all cases improves compared to transitional predictions based on known markers. As a result, we provide new opportunities to identify dissect lineage trajectories across transcriptionally disparate populations to make statistical predictions about gene expression networks underlying cell fate diversification.

Using polynomial regression analysis, we identified 483 genes whose expression levels significantly change across the differentiation time course, and comprised two discrete gene regulatory modules. These modules represent subsets of genes that either display high expression levels during the early stages of differentiation, or decreasing expression as the cells progress towards maturity, such as SOX2 and NANOG; or increasing expression as cells exit pluripotent stages towards maturing cardiac cells.

By integrating high-resolution single-cell transcriptomic data with population-based genomics data, we were able to identify cardiovascular diseases genes whose expression levels vary across the cardiac cell developmental lineage, implicating variation in the pathogenicity of GWAS loci in cell developmental lineages. This approach is especially relevant for cardiac diseases, as it has been demonstrated that cardiac reprogramming and differentiation preserve patient-specific expression patterns (Matsa, et al., 2016). Recent single cell studies have uncovered lineage-specific gene regulation programs that cause abnormal congenital heart defects (Delaughter et al., 2016). Here, we identified tropomyosin 1 (TPM1) as highly upregulated in D15S:S2 and D30:S2 subpopulations. TPM1 is crucial for muscle contraction and is also linked to several cardiac-related diseases, specifically familial hypertrophic cardiomyopathy, left ventricular non-compaction and familial dilated cardiomyopathy.

In addition to TPM1, we identified a number of important cardiac disease-related genes that play a significant role in cardiomyocyte differentiation and development. The AKR1B1 gene, which encodes aldose reductase was present in the ‘late’ module, and is also an eQTL regulated gene in adult left ventricle. In mouse models of ischaemia, there is evidence that inhibition of aldose reductase expression reduces ischemic injury (Ramasamy and Goldberg, 2010). Also present in the ‘late’ module were genes whose expression has been identified as playing a causal role in cardiovascular disease, including ADBR2 and MADD. SNPs within ADBR2 are significantly associated with cardiac disease, including atherosclerosis (Zak et al 2008), ischemic stroke (Stanzione et al, 2007), myocardial infarction and CAD (Wang et al, 2015). Loci in MADD are associated with coronary heart disease risk and ischemic stroke (Wu et al, 2016), as well as early diastolic heart failure (Wu et al, 2012). Given these genes are significantly associated with cardiomyocyte lineage specification, our results suggest that aberrant cardiac gene expression caused by regulatory SNPs during early cell fate determination may contribute to disease states that manifest later in life.

In summary, we have shown that high resolution single cell RNA-seq can be used to understand the complex cell fate decisions that occur during differentiation of iPSCs into mature cardiomyocytes, which is crucial for regenerative medicine and bioengineering. When combined with population-level genomic data, it can inform the function and potential pathogenicity of cell subpopulations during development.

## Methods

### Cell culture and cardiomyocyte differentiation

WTC CRISPRi human induced pluripotent stem cells (iPSC) were cultured on vitronectin coated plates in mTeSR media (Stem Cell Technologies, 05850) as previously described (Nguyen, *et al.,* under-review). Cultured iPSCs at a density of 3.5 x 10^5^ were differentiated into mature cardiomyocytes using a modified monolayer protocol described in detail in Friedman *et al.* (under-review). Briefly, cells were cultured in mTeSR media in 24-well plates at 37^o^C, 5% CO2. After 24 hours, cells (hereon day 0) were treated with a cocktail of 3μM CHIR-99021 (Stem Cell Technologies, 72054), 500μg/mL BSA (Sigma Aldrich, A9418), and 213μg/mL ascorbic acid (Sigma Aldrich, A8960) in RPMI (Life Technologies Australia, 11875119). At 72 hours, the media was replaced with RPMI containing 500μg/mL BSA, 213μg/mL ascorbic acid and 1μM Xav-939 (Stem Cell Technologies, 72674). After 120 hours, the media was replaced with RPMI containing 500μg/mL BSA and 213μg/mL ascorbic acid, and from 168 hours onwards, RPMI containing 1x B27 supplement plus insulin was replaced every 48 hours.

### Single cell isolation

Five time-points were selected for the duration of a differentiation time course. For each time point, 2 x 24-well plates of adherent cells were washed with PBS and harvested using a Versene + 0.25% Trypsin treatment, which was neutralized with 50% foetal bovine serum (Scientifix, FFBS-500) and 50% of DMEM/F12 media (Life Technologies Australia, 11320033). Pools of cells were generated using 12 wells of cells from each plate, spun at 300 g for 5 minutes and resuspended in Dulbecco’s PBS (Gibco; Cat#14190) with 0.04% bovine serum albumin (Sigma Aldrich, B6917). A BD Influx instrument was used to sort single, viable cells using Propidium Iodide into Dulbecco's PBS + 0.04 % bovine serum albumin, which were retained on ice. Sorted cells were counted and re-assessed for viability with Trypan Blue using a Countess automated counter (Invitrogen), and then resuspended at a concentration of 800-1000 cells/μL (8x10^5^-1x10^6^ cells/mL). Final cell viability estimates ranged between 87-95 %.

### Generation of single cell GEMs and sequencing libraries

Single cell RNA-Seq (scRNA-Seq) was performed in duplicate across a cardiomyocyte differentiation time course. High-throughput droplet partitioning of viable cells with barcoded beads was performed using the 10X Genomics Chromium instrument (10X Genomics) and the Single Cell 3′ Library, Gel Bead and Multiplex Kit (v1; 10X Genomics; PN-120233). The number of cells in each reaction was optimized to capture approximately 5,000 cells. cDNA was prepared from the resulting partitioned samples, and cDNA shearing was performed with a Covaris S2 instrument (Covaris) set to produce a target size of 200bp as per the manufacturer’s recommendation (Intensity:5, Duty cycle: 10%; Cycles: 200; Time: 120s). The resulting single cell transcriptome libraries were pooled and sequenced on an Illumina NextSeq500, using a 150-cycle High Output reagent kit (NextSeq500/550 v2; Illumina, FC-404-2002) in standalone mode as follows: 98bp (Read 1), 14bp (I7 Index), 8bp (I5 Index), and 10bp (Read 2).

### Bioinformatics mapping of reads to original genes and cells

Raw sequencing data (BCL) was processed directly with the *cellranger* pipeline v1.3.1 (*mkfastq*, *count*, *aggr*) using the default parameters, which were adjusted to provide an expected number of cells (5,000). The reads were aligned to the GRC38p7 human reference genome using the STAR software (Dobin et al., 2013) included in the *cellranger* pipeline. Quality control for cell barcodes and unique molecular identifiers (UMI) was performed using default parameters in the *cellranger count* processing. The final, between samples normalised expression matrix for 10 samples spanning the differentiation time course was generated using the *cellranger aggr* function.

### Interactive web server and data accessibility

RNA-seq data have been deposited in the ArrayExpress database at EMBL-EBI (www.ebi.ac.uk/arrayexpress) under accession number E-MTAB-6268. To facilitate the broad use of this single-cell human pluripotent single-cell dataset, we created an open, interactive R shiny server, available at http://computationalgenomics.com.au/shiny/hipsc2cm/. The server contains user-friendly data exploration and representation tools. Data can be interactively explored the expression of any gene in each cell of the 43,168 cells across five time points, and compared the expression of the selected genes between different subpopulations.

### Quality control

Filtering of cells was performed after aggregating samples, ensuring subsequent analysis was not affected by cells that were inconsistently outside the threshold of three times the Median Absolute Deviation (MAD):

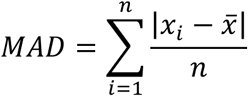

Where *x* is a vector of total mapped reads per cell, the total number of genes detected in a cell, or expression of mitochondrial and ribosomal genes. We filtered 87 cells with total mapped reads > 3 x MAD, and 300 cells with total number of genes > 3 x MAD. After removing cells, we subsequently removed 15,302 genes that were detected in less than 0.1% of all remaining cells.

### Normalisation

We normalised the expression data on three levels: between days, between samples, and between cells. To account for variation in total read depth per sample, we used a subsampling process based on the sample index, cell barcodes, and UMIs to randomly sample from a binomial distribution to the level of individual reads. This approach maintains the distribution of reads mapped per gene, per cell and per sample, and equalizes the total read-depth between samples. This method has been shown to be less biased than other global-scaling methods, which use information from the final mapped expression matrix only - *i.e.* total reads mapped per sample and distributions of expression levels in a sample (Zheng et al. 2016). For each sample (*i*) the subsampling rate (*Rate_i_*) was determined as:

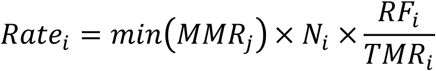

Where *MMR_j_* is the ratio of expected total reads divided by the expected mean reads per cell of all samples, with the minimum *MMR_j_* from all samples merged if fitted; *N_i_* is the number of cells in sample *i*; *RF_i_* is the fraction of mapped reads in sample *i* to the total number of mapped reads; and *TMR_i_* is the total number of mapped reads that share the same sample index.

For each gene in each cell, from the total set of reads that were mapped to the gene, a subset was randomly drawn from the binomial distribution, at the rate *Rate_i_*, 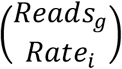. This process is more robust than standard-scaling approaches because it takes into account the unique read information associated to mapped genes and cells. Following resampling, the MMRs for the 10 samples were scaled, while the expression data distribution for genes in all cells of the sample was maintained.

After between sample normalisation, a deconvolution and pooling approach was performed to normalise read depth between cells. This method overcomes the inflation of zero count measures in the expression matrix of scRNA-seq data (otherwise known as dropout rate), by sequentially pooling 40, 60, 80 and 100 cells. The expression values of a gene across cells are summed to estimate a size factor for each pool. Pool size factors are then deconstructed into the size factors of individual cells. To estimate the scaling size factor for each cell, a deconvolution method (Lun et al., 2016) was applied for summation of gene expression in groups of cells. This summation approach reduces the number of stochastic zero expression of genes that are lowly expressed (higher dropout rates), or genes that are turned on/off in different subpopulations of cells.

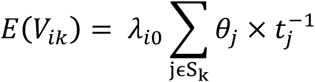

Where *S_k_* is a pool of cells, *V_ik_* is the sum of adjusted expression value (*z_i j_* = *θ_j_* × *λ_i_*_0_ where *λ_i_*_0_ is the expected transcript count and *θ_j_* is the cell specific bias) across all cells in pool *V_k_* for gene *i*, 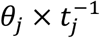 is the cell-specific scale factor for cell *j* (where *t_j_* is the constant adjustment factor for cell *j*).

The estimated size factor of a gene in a pool *S_k_*, termed as *E* (*R_ik_*), we calculate the ratio between the estimated *V_ik_* and the average *Z_i j_* across all cells in the population, such that

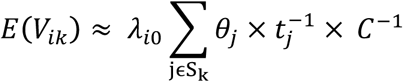

, with C defined as

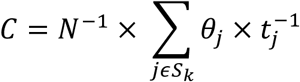

where *N* is the number of cells, *S*_0_ represents all cells and is a constant for the whole population and thus can be set to unity or ignored. The cell pools were sampled using a sliding window of cells ranked by library size for each cell. Four sliding windows with 20, 40, 60, and 80 cells were independently applied and results were combined to generate a linear system that can be decomposed by QR decomposition to estimate 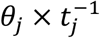 size factor for each of the cell. The final normalized counts are calculated by taking the raw counts divided by cell-specific normalized size factors.

### Dimensionality reduction

Normalized data were subjected to dimensionality reduction prior to clustering. Dimension reduction enables coordinates of a single cell to be represented in a two or three-dimensional space, allowing the relative location of single cell to be shown. We applied multiple independent approaches for representing and organizing cells in low dimensional spaces, including linear orthogonal transformation PCA (Principal Component Analysis), MDS (Multidimensional Scaling Principal Coordinate Analysis), non-linear methods (*t*-SNE and Monocle), and an imputation approach for circumventing the zero-inflation characteristic of a scRNA dataset (CIDR, as described in Lin et al., 2017). PCA and MDS are similar approaches that apply orthogonal transformation to the initial Euclidean distance matrix derived from the full expression matrix containing all genes and cells.

### Data visualisation in low dimensional space

We used Diffusion maps and t-SNEs for visualising gene expression and differentiation trajectory, but not for clustering or for inferring the final trajectories. We calculated the coordinates of each cell in *t*-SNE two (or three) dimensional map. Initially, ℝ^2^ were calculated based on a non-linear transformation of similarity between two cells derived from the initial simple Euclidean distance *d*(*C_i_*,*C_j_*) into *t*-SNE similarity matrix. The original pairwise similarity between two cells *C_i_* and *C_j_* in the original multidimensional space (ℝ^*G*^, G is the number of genes) was calculated as the joint probabilities 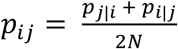, where *N* is the number of cells, and where the conditional probability of cell *C_j_* given cell *C_i_* was calculated as:

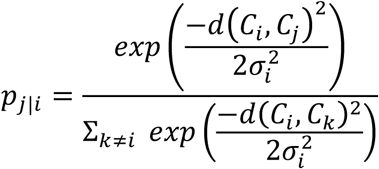

In which the *σ_i_* is the optimal bandwidth of the Gaussian kernel for cell *C_i_* such that the perplexity of the probability distribution for this cell in the low dimensional space equals a predefined constant perplexity. Thus, cells in the denser part of the data space have smaller *σ_i_*. In the low-dimensional *t*-SNE space, the pairwise similarity between two cells *q_ij_* is estimated in a gradient descent optimization process to minimize the Kullback-Leiber divergence of the distribution from Q (original space) to P (low-dimensional *t*-SNE space) in the following equation:

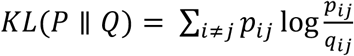

where *q_ij_* is the similarity score in the new space, calculated as

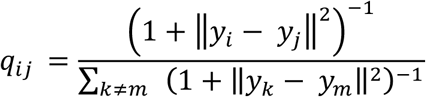

Thus, in the low dimensional space, the low similarity value between two cells suggests higher distance (*i.e.* cells are further away). The diffusion approach applies a diffusion distance matrix, which is an approximation of Euclidean distance matrix from a Gaussian sampling process to estimate a Gaussian center for each cell and a length scale that the cell can randomly diffuse (Haghverdi, et al., 2015). Pairwise transition probability between two cells *C_i_* and *C_j_* can be calculated as the interference of the two wave functions *Y_C__i_* (*x′ _i_*) and *Y_Cj_*.(*x′ _j_*). A *N* x *N* transition probability matrix can be calculated for all pairs of cells.

### Clustering at an Optimal REsolution (CORE)

We devised a novel unsupervised clustering method, which incorporates an approach to statistically identify the most stable resolution of clusters to identify distinct subpopulations of cells. We first construct an unsupervised dendrogram using the Euclidean distance matrix between cells, which is calculated using the normalised gene expression matrix (*n* cells x *p* genes). The Euclidean distance between two cells *i* and *j* can be represented as:

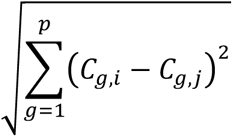

where *C_g,i_* is the expression of the gene *g* in the cell *C_i_*. Branching points in the dendrogram represent increasing smaller clusters of cells, with each branching point based on hierarchical clustering using the Ward method to minimise the within and between cluster variance (Murtagh and Legendre, 2014), calculated as the Euclidean distance to the centroid cell:

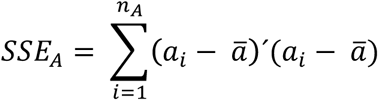

where 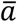 is the centroid cell of the cluster A, defined as the cell with the lowest sum of squares of all pairwise Euclidean distance within the cluster. The between cluster variance for the joint clusters {A, B} is calculated similarly. The hierarchical clustering begins with *n* clusters of size one. The two clusters with the minimal increase in the distance *SSE_AB_* − (*SSE_A_* + *SSE*_B_) are merged. The decision to merge the subsequent cluster (C) to the {A, B} requires C to satisfy the minimal value of two clusters, estimated by the Lance Williams algorithm (Lance and Williams, 1967):

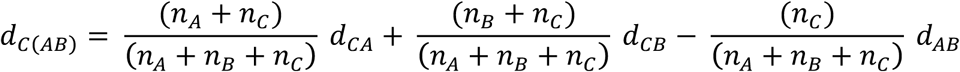

We subsequently apply a Dynamic Tree cut approach (Langfelder, et al., 2008) to merge clusters, based on the topology (shape) of the dendrogram. The Dynamic Tree cut is a top-down algorithm to firstly identify the largest clusters (based on a static height cut-off) and iteratively finds sub-clusters by analysing the fluctuation in the joining heights of the branches. The splitting is recursive until a stable number of clusters are identified. From the dendrogram, height for each branch is extracted into an ordered length vector *H* = (*h*_1_,*h*_2_, …,*h_n_*). The cluster merging process given a calibration height input *l* is based on the sign transition of the heights relative to *l*. The differences between each branch height and the calibration *l*, 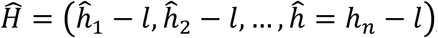, define turning points, where the 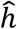 for consecutive branches in the ordered *H* turning signs to higher or lower than *l* The Dynamic Tree cut applies three calibration heights: *l_m_* (the mean calibration height 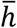), *l_u_* (the upper calibration height, 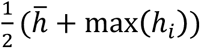, and *l_d_* (the lower calibration height, 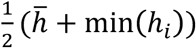 At each iteration, the three calibrationsheights *l_m_*, *l_u_* and *l_d_* are recalculated from *H*, thus making the procedure dynamic.

The Dynamic Tree cut will result in different numbers of clusters depending on the height threshold (denoted as *W_i_* described below) specified for merging branches. Furthermore, the dynamic merging process does not take into accounts the individual members (single cells) in each merged clusters. To solve the limitations we developed an approach to find an optimal clustering resolution (height threshold). The lower the height thresholds represent a higher resolution resulting in more clusters, while a larger height threshold leads to lower resolution, but fewer and more stable clusters. Our algorithm loops through the dendrogram, applying the Dynamic Tree cut, and comparing results between two consecutive steps during optimisation of the clustering based on the adjusted Rand index. The Rand index (Rand, W., 1971) measures the similarity between two clustering results based on the pairwise shared membership of any two cells. The Rand index is based on the assumptions that every cell within a cluster has equal weight in defining a cluster, two categories of membership (belonging or not belonging to a cluster) are equally important, and every cell is discretely assigned to one cluster. The adjusted Rand index implements a model for random assignment of points into cluster (Hubert and Arabie, 1985). The adjusted Rand index for two clustering results *C* and *C′,* containing *K* and *K′* clusters, are calculated by the equation below:

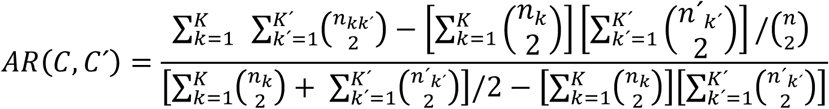

where *n_k_* and *n′_k_* are the cell numbers in clusters *C_K_* and *C′_K′_* of the two clustering results *C* and *C′*. *n_kk′_* = ∣*C_k_* ∩ *C_k′_*∣ is the number of cells in the intersection of clusters *C_K_* and *C′_K′_*.

Using these methods we select for the optimal cluster resolution by implementing the following algorithm:

1. Apply cutreeDynamic 40 times to merge branches in 40 different height windows (defined the dendrogram area to be merged) from bottom (*W*_1_ = [0.025,1]) to the top (*W*_1_ = [1,1]).
2. Compute pairwise adjusted Rand index (*AR_i_*) for every 2 consecutive windows (*W_i_* and *W_i+1_* for integers *i ∊* [1,39])
3. Compute stability index *S* spanning the 40 iterations. *S* is the set of count values *C_s_* for unique Rand index values *AR_i_* that remain the same between consecutive *W_i_*.
4. Determine the most stable clustering result *C_s_*, where *s* is selected by the following criteria:
*AR_S_* = max(*S*) and max(*S*) is different to *AR_1_* or *AR*_40_ *s* = 1 or 40 if *AR*_1_ or *AR*_40_ = max(S) and *C_s_* /40 >0.5(i.e. stable in more than 50% of all iterations)

### Cell cycle analysis

To assess whether the clustering assignments were influenced by the differences in cell cycle phases, we applied a machine-learning model to predict cell cycle phases (Scialdone et al., 2015; Lun et al., 2016). The model uses scRNA gene expression data, and a reference training set (prior-knowledge) of relative expression of “marker pairs”, in which the sign of each pair changes between cell cycle phases (Scialdone et al., 2015). We used a gene training set (containing ordered gene list for above 20,000 genes) from Leng et al. (2015). Scores for each of the three phases G1, G2M and S were estimated based on the proportion of the training pairs with sign changes in each phase relative to the other phases.

### Differential expression analysis

We performed differential expression analysis between subpopulations of cells identified independently for each time-point to address two scenarios: 1. The differences in gene expression between cells in one subpopulation compared to all remaining cells in a given time-point; 2. Differences between cells in one subpopulation and cells in a subpopulation in the subsequent time-point. Normalized gene expression level, were transformed from log2 value to read counts, and used as input into DESeq (Anders and Huber, 2010), with the following modifications. One pseudocount was added to the normalised expression values, before rounding, to generate a mapped read count matrix without Zero expression values. Due to the low expression values (relative to 1 as the added pseudocount), the fold-change values were readjusted post differential expression analysis by subtracting to 1. The *p*-values were adjusted by the Bonferroni correction (*p*-values multiplied by the total number tests), and significant genes are those with *p*-adjusted values lower than 0.05.

### Prediction of differentiation trajectory using a novel machine learning approach

We developed an unsupervised, machine learning approach for identifying differentiation trajectory and and cell fate. The method, which we term scGPS, includes two main steps. First, we construct and train the model using variable selection. For cells in a subpopulation, we select their *p* differentially expressed genes and apply a LASSO (Least Absolute Shrinkage and Selection Operators) regression to identify gene predictors. The *p* genes can be considered as markers for the subpopulation, distinguishing them from remaining cells. We classified the dataset into two classes, including *n* cells in a subpopulation of interest (class 1), and remaining cells that do not belong to class 1 (class 2). The *n/2* cells are removed from the original dataset, so that they represent 50% of the total cells for each of the two classes. Let cell-subtypes be a qualitative response variable *y* and assigns *y* into a class *k*, we estimate for each of the *n/2* cells belonging to one of the *k* classes. We construct a response matrix Y with element *Y_ik_* = *I*(*Y_i_ = k*), *k* ∊ (0,1) are two classes (belong or not belong to the subpopulation). Let *p* equal the number of gene predictors. Denote the conditional class probability of a cell *C_i_* belonging to a class *k* (0 or 1 belong or not belong to) given the gene expression profile *x_i_* of *p* genes *x_i_* = (*x*_1_,*x*_2_, …,*X_ip_*) as: Pr(*y* = *k* ∣*x_i_*). We then fit a generalized linear model (binomial distribution) with the response variable as a vector containing two classes, and the predictor as the matrix *n* by *p* of the expression levels for the class 1 cells Effects of the genes are estimated by a penalized maximum likelihood procedure. The conditional class probabilities of cell *C_i_* belonging to class *k* is the linear combination of selected genes:

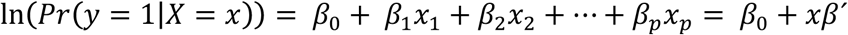

where *β_j_* is a coefficient for gene *j* (*β_j_* = 0 if the gene *j* is not a predictor of the class). The coefficient vector *β* = (*β*_0_,*β*_1_,*β*_2_,…,*β_P_*) are calculated by maximum likelihood estimation. The predicted probability of a cell *C_i_* being in a subpopulation 1 or 0 is estimated by replacing *β* and gene expression values to the regression equation (3). For each subpopulation, the resulting model with the optimal set of non-zero coefficient genes is a Bayes optimal classifier. The model removed insignificant genes *j* that do not contribute to the model fit by shrinking their coefficients to 0 following:

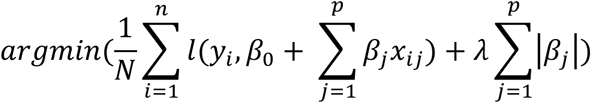

where *x_i_* = (*x*_1_,*x*_2_, …,*X_ip_*) is a vector of expression values of *p* genes in cell *C_i_*; *y_i_* is the cell subpopulation of the cell 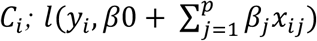 is the negative log-likelihood for *C_i_*; and *λ* is a tuning parameter that controls the shrinkage penalty of the coefficients. For each training cell subpopulation, an optimal *λ* and a set of gene predictors can be determined by a 10-fold cross-validation procedure to select the *λ* that produced the minimum classification errors. The LASSO procedure optimizes the combination set of coefficients for all predictors in a way that the residual sum of squares is smallest for a given *λ* value [38].

The final step is to estimate the prediction accuracy using 100 bootstrap results for each prediction. The optimal LASSO model with the highest prediction accuracy and the deviance explained is selected from the bootstrap replicates. The logistic regression model containing the optimal set of LASSO selected genes and corresponding coefficients are applied for prediction of the percent of cells in a targeted (data not in the training set) subpopulation.

### Identifying significance changes in gene expression across the cardiomyocyte differential lineage

We applied a polynomial linear model framework to identify genes whose expression levels changed over the actual differentiation time post induction. We fit the expression levels of each gene across five time-points into a cubic regression model. Prior to the modelling step, the expression data in mapped reads per gene, was transformed into Z-score such that each gene had the mean expression across all cells equal 0 and standard deviation equal 1. We transformed the categorical variable representing one of the five time-points to a quantitative scale, with equal weights for each time-point. Changes in the expression levels across the differential time course were tested using the following function:

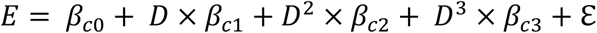

Where *E* is the matrix of expression value for 17,718 genes across all 43,168 cells assigned into five time-points D = (1, 2, 3, 4, 5) and *β_c_*_0_,*β_c_*_1_,*β_c_*_2_ and *β_c_*_3_ are vectors of coefficients for each gene in the cubic models respectively. For each model, p-values, regression coefficients, coefficient of determination (R-squared) and adjusted R-squared (adjusted for the number of predictors used in the model) were calculated. The adjusted R-squared values were used to assess the proportion of gene expression variance explained by the cubic terms. Significant (*p* < 0.05/17,718) genes were further analyzed by applying weighted co-regulation network analysis (WGCNA) to identify submodules of co-regulated genes with significant association to time. Genes in each submodule were analyzed for enrichment of transcription factor motif by iRegulon and Reactome pathways.

### Identifying cardiovascular disease risk variants acting though changes in cardiac tissue gene expression

To identify genes that have a pathogenic role in cardiovascular disease, we integrated population scale GWAS and eQTL data from cardiac tissue and used a Summary Mendelian Randomization (SMR) method to identify shared causal loci. GWAS summary statistics for 4 analyses from the Cardiogram study (Cardiogram, CardiogramPlusC4D, C4D CAD Discovery and MICAD) (Schunkert et al (2011), Coronary Artery Disease (C4D) Genetics Consortium (2011), Webb et al (2017)) were combined to yield a total of 2,962,408 (2,447,611 unique) SNPs with MAF >= 0.01. Expression Quantitative Trait Loci (eQTL) data was obtained for human cardiac tissue-specific via the Genotype-Tissue Expression (GTEx) project for Left Ventricle (LV) and Atrial Appendage (AA), and Left Ventricle from Koopmann *et al* (2014), and individual-level genotypes and peripheral blood eQTL data of 2,765 individuals were obtained from the CAGE study (Lloyd-Jones et al, 2017). SMR software (Zhu *et al.* 2016) version 0.67 was used to prioritise genetic variants with highly significant causal effects. Full details of the Summary Mendelian Randomization (SMR) method can be found in the Zhu *et al.* (2016). However, we used SMR to jointly analyze cardiovascular disease GWAS and cardiac tissue eQTL summary statistics to test if there is a shared causal variant at the locus for both the disease and gene expression. SMR uses SNPs (Z) to test whether an exposure (X) has a causal effect on an outcome (Y), where Y is cardiovascular disease and X an eQTL (VanderWeele TJ Epidemiology 2014; Boef Int J Epidemiology 2015). SMR estimates the effect of X on Y (^b^_XY_) as ^b^_XY_ = ^b^_ZY_^/b^_ZX_, where ^b^_ZY_ is the effect size of Z on Y and ^b^_ZX_ is the effect size of Z on X (Zhu et al., 2016). To test the hypothesis that the GWAS hits for cardiovascular disease and cardiac tissue eQTL (X) have the same causal loci, SMR estimates the SNP effect (^b^_ZY_) from GWAS summary data, and the SNP effect on gene expression (^b^_ZX_) from summary data of eQTLs. SMR tests for the following models; causality (Z → X → Y), pleiotropy (Z → X and Z → Y), and linkage (Z_1_ → X, Z_2_ → Y, and Z_1_ and Z_2_ are in LD). The model of linkage is excluded using the heterogeneity in dependent instruments (HEIDI) test, which considers the pattern of associations using all the SNPs that are significantly associated with eQTLs in the *cis*-region. The HEIDI test takes into account non-independence of *cis*-eQTLs due to LD (using individual-level data from a reference sample to estimate LD between the *cis*-SNPs). Genes that show evidence of heterogeneity (e.g. *p*_HEIDI <0.05) are rejected. The null hypothesis is that there is a single causal variant affecting and cardiovascular disease risk and gene expression (pleiotropy or causality). The alternative hypothesis is that gene expression and cardiovascular disease risk are affected by two distinct causal variants. The genes that had a causal variant with a highly significant *p*_SMR value and *p*_HEIDI > 0.05 were retained.

## Acknowledgements

Sequencing was performed by the Institute for Molecular Bioscience Sequencing Facility at the University of Queensland. The WTC CRISPRi hiPSCs and pQM plasmid backbone were kindly provided by the Conklin lab (UCSF, Gladstone Institute). This work was supported by the Australian Research Council SR1101002 (NJP), and National Health and Medical Research Council Fellowship 1107599 and grant 1083405.

## Lists of Supplementary Figures

**Figure S1**. Overview of the analysis pipeline for the 44,123 cells at the five stages of cardiomyocyte differentiation.

**Figure S2.** Single cell RNA sequencing results of 44,123 (before filtering) in 10 samples with 2 biological replicates for each of the 5 time points.

**Figure S3.** Overview of the merged dataset by days, biological replicates and general clusters.

**Figure S4.** The clustering at optimal resolution (CORE) method.

**Figure S5.** Known markers confirmed separation of time-points by differentiation stages.

**Figure S6.** Gene markers for subpopulation at day 5.

**Figure S7.** Gene markers for subpopulations in day 2, day 15, and day 30.

**Figure S8.** A publically available resource with user-friendly tools to mine the large scale singlecell transcriptomics dataset.

**Figure S9.** Cell cycle analysis of all cells in all clusters and days.

**Figure S10.** Transition scores between subpopulations estimated by GPS.

**Figure S11.** Prediction accuracy by GPS models for each of the subpopulation.

**Figure S12.** Differentiation pseudotime analysis by Monocle and Diffusion methods. Analysis

**Figure S13.** Regulatory enrichment of gene targets for significant genes identified by cubic modelling.

## List of Supplementary Tables

**Table S1.** General quality control statistics for each of the 10 sequenced samples.

**Table S2.** Summary of the data filtering process to remove outlier cells and genes.

**Table S3.** Mean expression of 140 known pluripotency and cardiomyocyte differentiation markers.

**Table S4.** Differentially expressed genes and pathways between two continuous days.

**Table S5.** Differentially expressed genes distinguishing each of the subpopulations.

**Table S6.** Cell cycle phase prediction for each subpopulation.

**Table S7.** Subpopulation transition scores between and within time-points (mean +- standard deviation for 100 bootstrap iterations)

**Table S8.** LASSO selected genes and coefficients.

**Table S9.** Summary statistics for significant genes in linear, quadratic and cubic models.

**Table S10.** Cubic genes and pathways early and late modules

**Table S11.** Transcription factor binding target enrichment analysis for significant cubic genes in early and late modules, and in trajectory 1 and trajectory 2 genes.

**Table S12.** Significant genes and pathways in the cubic module for cells in independent trajectory 1 and trajectory 2.

**Table S13.** Overlapping genes between significant cubic genes from single cell analysis and eQTL genes from Koopman et al. (2014)

**Table S14.** Overlapping genes between SMR and single cell analysis gene lists (significant cubic genes, differentially expressed genes and LASSO predictor shrinkage genes)

**Table S15.** Mann-Whitney U test for difference in gene expression of 21 CAD genes across five time-points

**Table S16.** Expression of 101 CAD genes reported in Howson et al. (2017)

